# Environmental conditions dictate differential evolution of vancomycin resistance in *Staphylococcus aureus*

**DOI:** 10.1101/2020.06.07.138933

**Authors:** Henrique Machado, Yara Seif, George Sakoulas, Connor A. Olson, Richard Szubin, Bernhard O. Palsson, Victor Nizet, Adam M. Feist

## Abstract

While microbiological resistance to vancomycin in *Staphylococcus aureus* is rare, clinical vancomycin treatment failures are common, and methicillin-resistant *S. aureus* (MRSA) strains isolated from patients after prolonged vancomycin treatment failure remain susceptible. Adaptive laboratory evolution was utilized to uncover mutational mechanisms associated with MRSA vancomycin resistance in a bacteriological medium used in clinical susceptibility testing and a physiological medium. Sequencing of resistant clones revealed shared and media-specific mutational outcomes, with an overlap in cell wall regulons (*walKRyycHI, vraSRT*). Evolved strains displayed similar genetic and phenotypic traits to resistant clinical isolates. Importantly, resistant phenotypes that developed in physiological media did not translate into resistance in bacteriological media. Further, a bacteriological media-specific mechanism for vancomycin resistance enabled by a mutated *mprF* was confirmed. This study bridges the gap of understanding between clinical and microbiological vancomycin resistance in *S. aureus* and expands the number of allelic variants that result in vancomycin resistance phenotypes.

## Introduction

Antibiotic resistance is a global healthcare threat worldwide (Organization, 2014; (u.s.) and Centers for Disease Control and Prevention (U.S.), 2019). Consequently, numerous strategies have been developed and implemented to monitor, assess, and circumvent the development of antibiotic resistance among pathogens (Alcock et al., 2020; Martens and Demain, 2017; Pollack and Srinivasan, 2014). Continual monitoring and assessment are key to getting a global picture of the problem and increasing our understanding of the mutational mechanisms that pathogens employ towards resistance development.

Although somewhat successful, current monitoring and assessment approaches are based on existing pathogen-specific knowledge. Because mechanisms for antibiotic resistance evolution are poorly defined, full realization of critical threats often occurs only after resistance has emerged. Importantly, the evaluation of allelic variations known to lead to reduced susceptibility to a given antibiotic do not account for all the other variations of that same allele or variation in other alleles that can result in the same phenotype (Mitsakakis et al., 2018). Furthermore, bacterial susceptibility to antibiotics is measured following guidelines of the Clinical & Laboratory Standards Institute (CLSI), which recommend using the bacteriological rich media cation-adjusted Mueller-Hinton broth (CA-MHB) to determine antibiotic susceptibility. CA-MHB was specifically developed for its ability to reliably support the cultivation of common human pathogens from clinical samples, and only later adopted for minimum inhibitory/bactericidal concentration (MIC/MBC) testing of antibiotic candidates. However, CA-MHB does not come close to recapitulating the environment encountered by bacteria *in vivo* and has been shown to be less reliable in predicting *in vivo* activity of antibiotics than other more physiological media such as mammalian tissue culture media (Ersoy et al., 2017; Farha et al., 2018; Kumaraswamy et al., 2016). Adaptive laboratory evolution (ALE) is a strategy that allows the investigator to address both the issues of limited coverage of allelic variation and environment-specific susceptibility through the study of identification of causal mutational mechanisms(Dragosits and Mattanovich, 2013; Mohamed et al., 2017; Salazar et al., 2020). ALE leverages microbial growth under different environments and conditions, wherein the natural mutation rate of bacteria can be exploited to sample successful allelic variations.

*Staphylococcus aureus* is a historical example of the successful development of antibiotic resistance by a common human pathogen, with methicillin-resistant strains (MRSA) presenting significant treatment challenges (Howden et al., 2010; Mwangi et al., 2007; Pader et al., 2016). The most commonly recommended drug for the treatment of MRSA infections is the glycopeptide vancomycin (Sorrell et al., 1982). Even though very few vancomycin-resistant MRSA clinical isolates have been reported (Hidayat et al., 2006; Hiramatsu et al., 1997; Howe et al., 1998), an increasing challenge of clinical treatment failures is well documented (Howden et al., 2010). The established MIC breakpoints determined by CLSI classify *S. aureus* into three susceptibility categories: vancomycin-susceptible *S. aureus* (VSSA, MIC ≤ 2 μg/mL), vancomycin intermediate-resistant *S. aureus* (VISA, MIC = 4-8 μg/mL), and vancomycin-resistant *S. aureus* (VRSA, MIC ≥ 16 μg/mL).

Here we used a clinical MRSA isolate (TCH1516) and applied adaptive laboratory evolution to uncover mutational mechanisms associated with resistance under two different environmental conditions: (i) CA-MHB, the nutrient rich bacteriological medium used for clinical susceptibility testing by CLSI recommendations; and (ii) Roswell Park Memorial Institute medium (RPMI), a mammalian cell culture medium that better mimics human physiology (McKee and Komarova, 2017). We further phenotypically characterized several vancomycin-tolerant and -resistant clones and identified genetic mutations responsible for such adaptations. The mutational evolutionary pathways towards vancomycin tolerance exhibited media specificity, with an overlap in regulatory rearrangements in cell wall regulons. We also establish that vancomycin-resistant phenotypes that developed in physiological media do not translate into resistance in bacteriological media, where a major resistance mechanism relies on change of the cell surface charge by mutation of *mprF*. This study significantly expands knowledge of allelic variation that contributes to *S. aureus* vancomycin tolerance.

## Results

### Tolerization of *S. aureus* TCH1516 to vancomycin

Adaptive Laboratory Evolution (ALE) relies on the natural capability of cells to adapt to new environments. Here, we have applied this technology to engender tolerance of *S. aureus* TCH1516 to vancomycin, and unravel the molecular mechanisms for this adaptation, in a so-called Tolerization ALE (TALE) (Mohamed et al., 2017; Salazar et al., 2020). Two media types were used in this experiment: CA-MHB, an undefined defined nutrient rich bacteriological medium used for clinical susceptibility testing and the well-defined Roswell Park Memorial Institute medium (RPMI), routinely used in the culturing of mammalian cells, and which resembles the physiological conditions in the human body (McKee and Komarova, 2017), supplemented with 10% bacteriological rich Luria-Bertani medium (RPMI+) to ensure bacterial growth equivalency. For each media type, four replicates of wild-type (WT) and of two media-adapted clones were evolved to stepwise increasing concentrations of vancomycin (Figure 1A). The TALE experiments were conducted for ∼30 days and ∼5×10^12^ cumulative cell divisions (CCDs). The final vancomycin concentrations reached an average of 5.14 ± 0.46 μg/mL in CA-MHB and 6.13 ± 1.03 μg/mL in RPMI+ (Figure 1B), compared to a tenth of the MIC used as start concentration (MIC in Supplementary Table 1; starting concentration of 0.1 and 0.2 μg/mL, in CA-MHB and RPMI+ TALEs, respectively).

**Figure 1.**
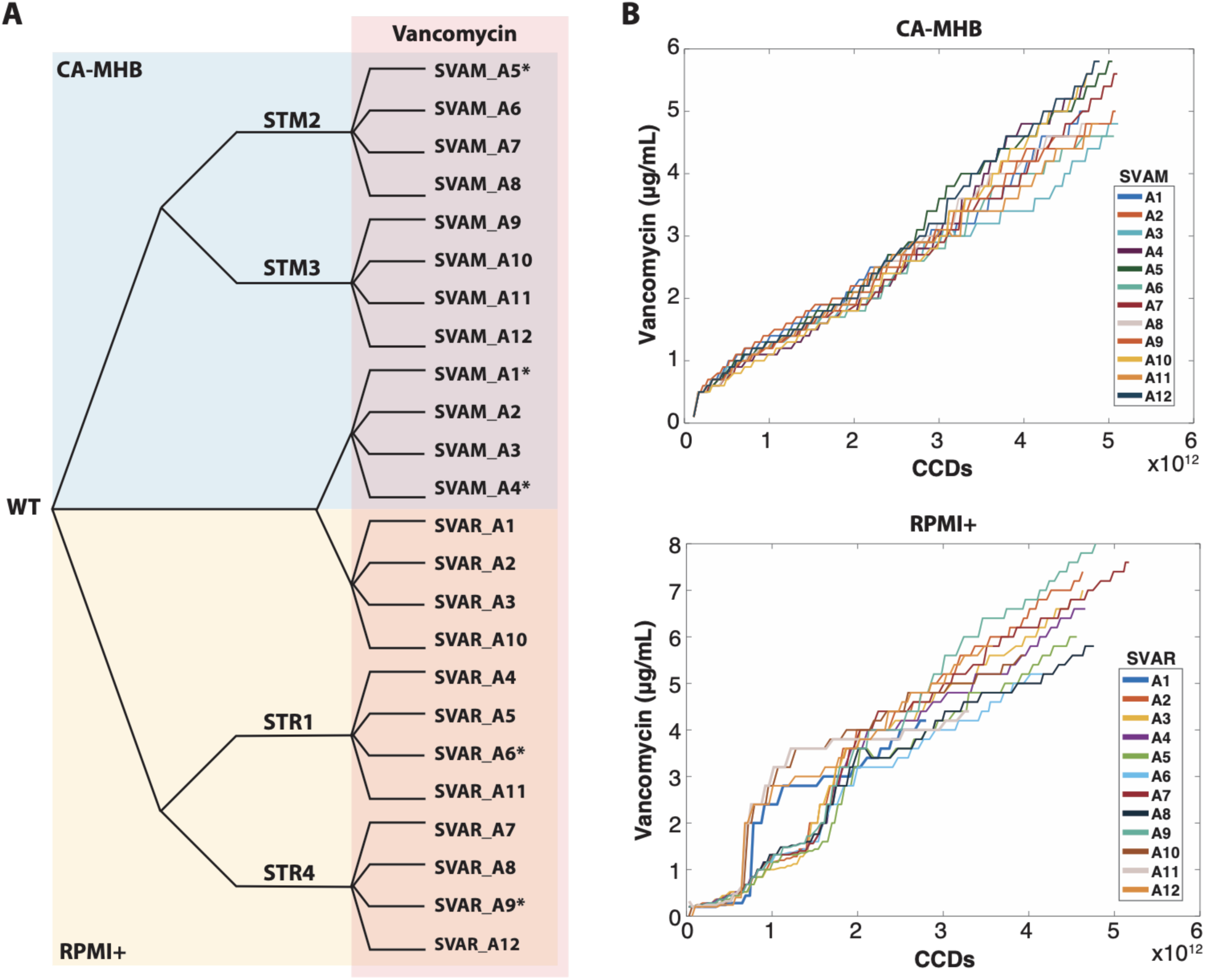
Tolerization Adaptive Laboratory Evolution. (A) Experimental design. Wild-type (WT) *S. aureus* TCH1516 was used as starting strain along with strains which were initially adapted to each media condition (Salazar et al., 2020). Strains were tolerized under CA-MHB (blue) and RPMI+ (yellow) media conditions with increasing concentrations of vancomycin (red). Isolate naming from each lineage is listed. (B) Plots showing the stepwise increase of vancomycin throughout the TALE experiments. * denotes hyper-mutators. STM: *Staphylococcus aureus* adapted to CA-MHB. STR: *Staphylococcus aureus* adapted to RPMI+. SVAM: *Staphylococcus aureus* tolerized to vancomycin in CA-MHB. SVAR: *Staphylococcus aureus* tolerized to vancomycin in RPMI+. CCDs: cumulative cell divisions.

### Phenotypes and tradeoffs in vancomycin tolerization

The TALE evolved strains, adapted for growth in increasing concentrations of vancomycin, were evaluated for changes to their growth phenotypes and antibiotic susceptibility. Growth rate was not affected in TALE strains, but an increase in the lag phase could be identified in CA-MHB-evolved strains when measured with no vancomycin stress and compared to their pre-evolved counterparts (Figure 2A). As expected, TALE strains grew in higher concentrations of vancomycin, with an increase in MIC of up to 8-fold (Supplementary Table 1). Previous studies have outlined the phenotypic characteristics of clinically isolated vancomycin-tolerant strains (Howden et al., 2010; Ishii et al., 2015). We observed similar characteristics in the TALE-evolved strains, including lower hemolytic activity and reduced autolysis (Supplementary Figure 1A and 1B). Further, vancomycin-tolerant strains generated in the laboratory have been reported to be phenotypically unstable (Gardete et al., 2012), losing their tolerance after growth in non-selective conditions. Therefore, we grew the vancomycin-tolerized strains for 21.79 ± 2.08 passages (9.41×10^11^ ± 9.84×10^10^ CCDs) in the media used for evolution, without vancomycin. The endpoint strains were generally stable in maintaining their tolerance phenotype in 11 out of 12 lineages, with the exception of the SVAM_A10 lineage, which decreased its MIC from 8 to 2 μg/mL (Supplementary Table 1).

**Figure 2.**
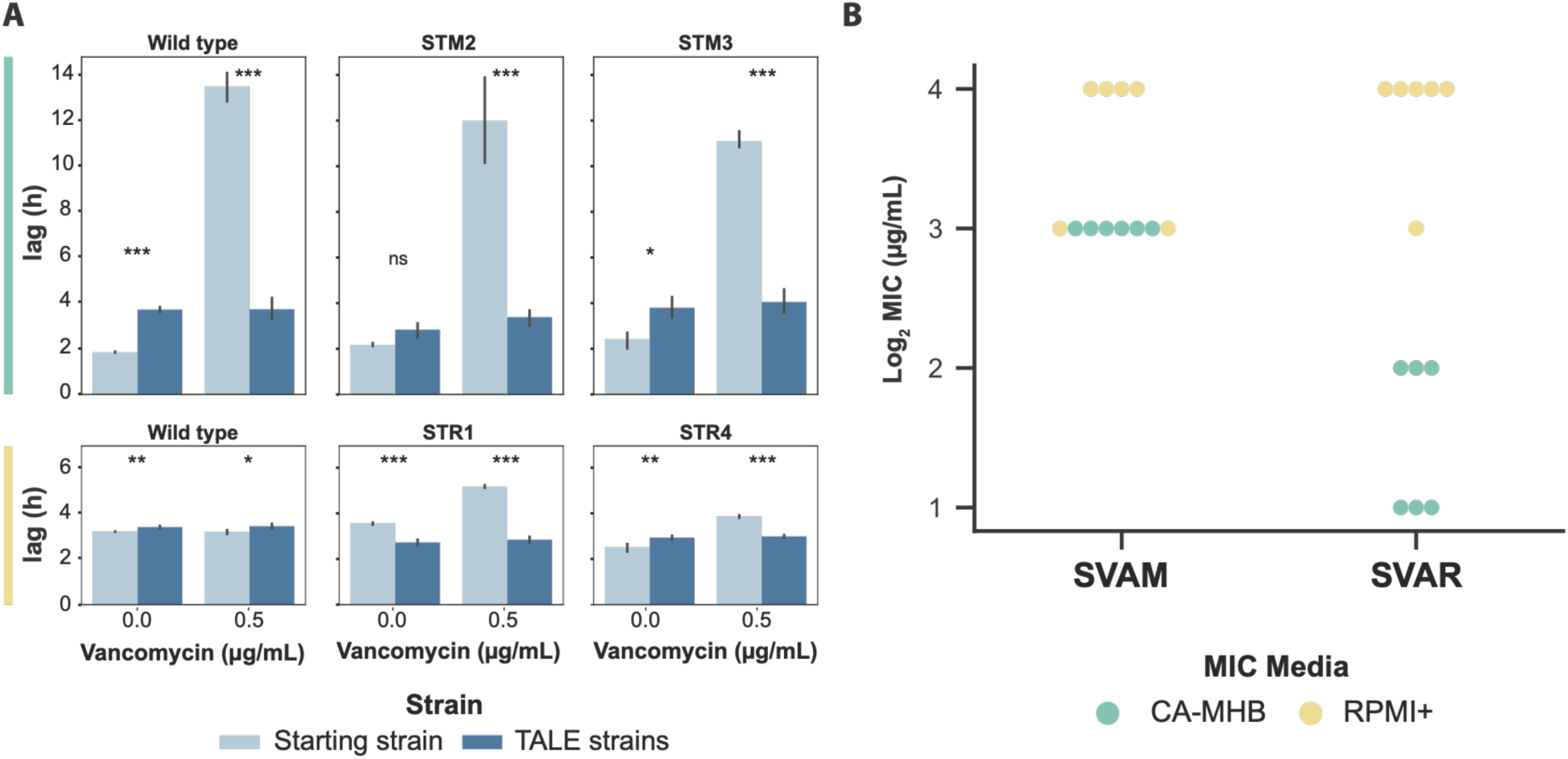
Phenotypic changes observed in vancomycin-tolerized strains. (A) Bar plots displaying growth trade offs grouped by the starting strain for each set of TALE experiments. Evolution starting strains and vancomycin adapted strains were grown in the absence and presence of vancomycin. Assessment of the lag-phase duration of starting strains and evolved strains in the same media as that used for tolerization, CA-MHB (top, green) and RPMI+ (bottom, yellow). Evolved strain averages used 2, 3 or 4 distinct clones derived from the given starting strain (Supplementary Table 1), all determinations were made in triplicate. Values that are significantly different by ANOVA are indicated by asterisks (ns, non-significant; *, P ≤ 0.05; **, P ≤ 0.01; and ***, P ≤ 0.001) (Supplementary Table 2). (B) A plot of Log_2_ vancomycin MIC values of evolved strains from all TALE conditions tested in both media types. SVAR strain tolerance phenotypes did not translate when tested for MIC in CA-MHB media. SVAM: *Staphylococcus aureus* tolerized to vancomycin in CA-MHB. SVAR: *Staphylococcus aureus* tolerized to vancomycin in RPMI+.

TALE strains lost vancomycin susceptibility, with a corresponding increase in their MIC (Supplementary Table 1). We further assessed if this decrease in susceptibility held true across the different media environments. Although CA-MHB vancomycin-tolerized strains (i.e., SVAM) maintained their tolerance phenotypes in RPMI+, the same was not true for the clones evolved in RPMI+. RPMI+ vancomycin-tolerized strains (i.e., SVAR) did not show decreased susceptibility when screened in CA-MHB media (Figure 2B). Thus, the strains evolved under RPMI+ displayed a media-specific tolerance phenotype, as compared to the translatable phenotype of the CA-MHB derived strains, for the two media conditions tested here.

### Environment dependent mutational strategies to vancomycin tolerization

For each of the 24 independent adaptive evolutionary lineages, 2-3 clones were randomly selected at different time points of the TALE experiments and were sequenced for mutational analysis. For each media type, approximately 400 unique mutations could be identified (462 in CA-MHB and 374 in RPMI+, n=50) (Supplementary Tables 3 and 4), with the majority (approx. 85%) being single nucleotide polymorphisms (SNPs). The percentage of transitions and transversions was quite similar (55% transitions, 45% transversions), but there was a bias in mutations from GC to AT, which constituted about 40% of the SNPs compared to 20% from AT to GC. The observed AT biased mutation has been shown to be universal for bacteria, independently of their genomic GC content (Hershberg, 2015; Hershberg and Petrov, 2010; Hildebrand et al., 2010).

Key mutated genes were considered to be those mutated in two or more independent TALE lineages and that were present in at least one clonal sample. If a gene was mutated in multiple flasks of the same TALE lineage or was only observed in sequenced population samples, it was not considered. A total of 69 key mutations were identified for both media types (Supplementary Figure 2). Overall, clones evolved in RPMI+ typically had a lower number of mutations (7.4 ± 3.4 mutations per strain) compared to the ones evolved in CA-MHB (10.6 ± 4.6 mutations per strain), excluding hyper-mutators. This mutational count difference is also reflected in the number of key mutated genes identified in both conditions, 54 versus 26 in CA-MHB and RPMI+, respectively (Figure 3A). By increasing the threshold of lineages with a given gene mutated, the decrease in the number of key mutated genes in CA-MHB was striking, whereas in RPMI+ there was less variance (Figure 3A). Although the *apt* gene appears as a key mutation in RPMI+ because it was mutated in all four replicates that were started from wild-type, this mutation has been associated with a growth rate increase in RPMI+ and does not have an effect on antibiotic susceptibility (Salazar et al., 2020). Furthermore, mutations in *mutL* were found in TALE strains from both media conditions and similarly, these strains displayed hypermutator phenotypes similar to previous reports (Ban and Yang, 1998; Glickman and Radman, 1980). The hypermutator strains had a higher number of mutations, compared to other TALE strains. However, the mutations identified were not distinctly linked with vancomycin tolerance but randomly spread throughout the genome (Supplementary Tables 3 and 4).

**Figure 3.**
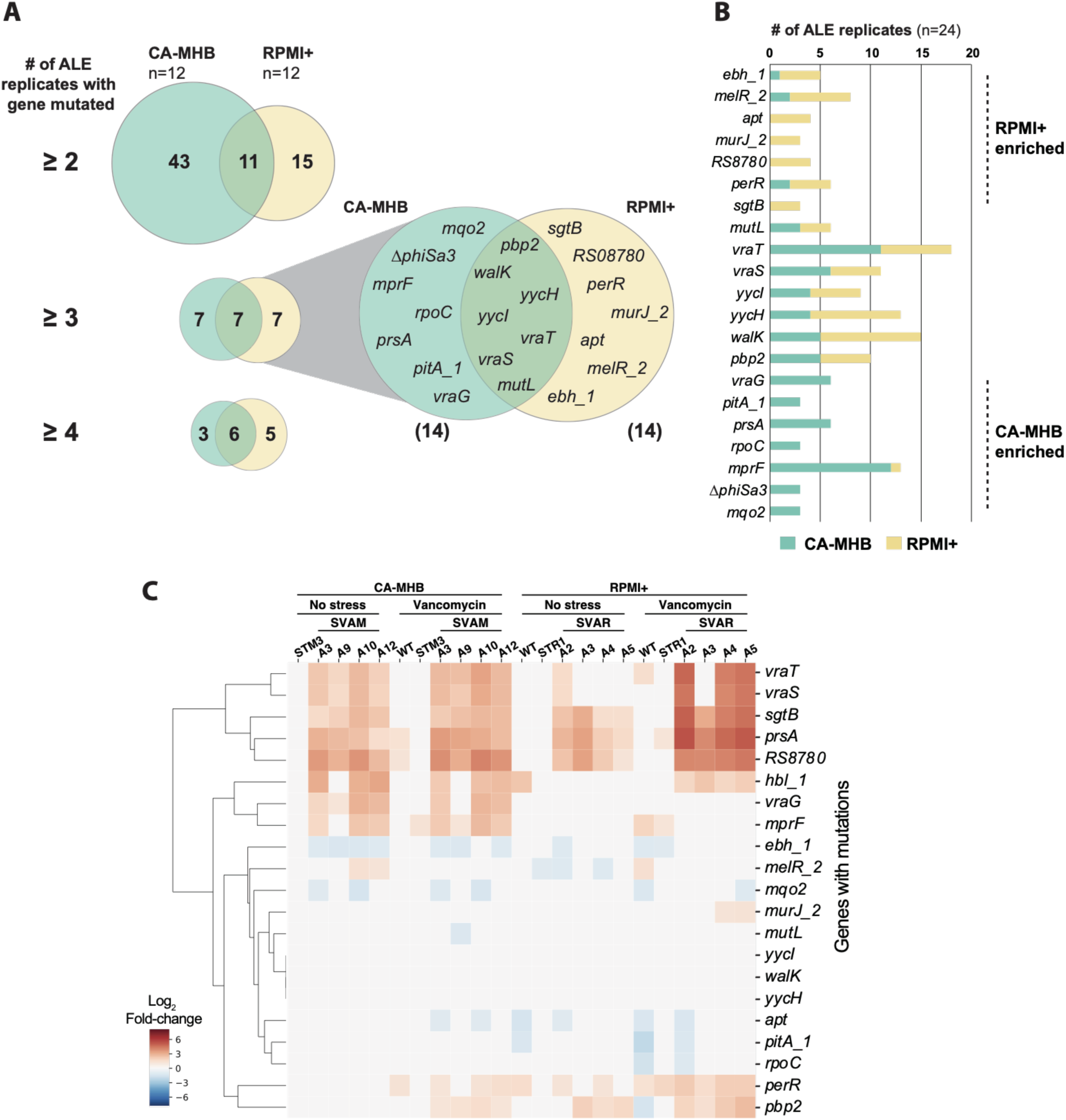
Key mutations found in MRSA during vancomycin tolerization. (A) Venn diagrams of the number of key mutated genes in the two utilized media conditions (i.e., CA-MHB and RPMI+). (B) A bar plot of the number of lineages with mutations in a key mutated gene, n=24 lineages, 12 for each media condition. (C) A heatmap of expression level of key mutated genes in a selection of starting and TALE-derived strains, in the presence and absence of vancomycin. STM: *Staphylococcus aureus* adapted to CA-MHB. STR: *Staphylococcus aureus* adapted to RPMI+. SVAM: *Staphylococcus aureus* tolerized to vancomycin in CA-MHB. SVAR: *Staphylococcus aureus* tolerized to vancomycin in RPMI+.

Distinct key mutations were identified in both bacteriological and physiological media conditions, with the overlap of similar mutations being mostly in regulatory genes, specifically in the *vra* and *wal* regulatory systems (Figure 3A). Mutations in these two systems have been previously associated with decreased susceptibility to glycopeptides (Gardete et al., 2012; Howden et al., 2010; Hu et al., 2016; Kato et al., 2010). For the *vra* system, comprised of the genes *vraSRT*, we identified 37 different mutations, of which 5 exact mutations (VraT-A151T, VraS-A314V, VraS-G88D, VraS-T264A, and VraR-V14I) and 2 mutated positions (VraT-P126 and VraT-N74) have been previously described (Cameron et al., 2012; Hu et al., 2016; Kato et al., 2010). For the *wal* system, comprising genes *walKRyycHI*, we identified 44 mutations, 26 of which were in the accessory genes *yycHI* (21 resulting in possible pseudogenization, i.e. gene disruption). This pseudogenization type of gene disruption (specifically, a frameshift mutation resulting in truncation) had been previously observed in an *in vivo* evolution study in a patient (Mwangi et al., 2007). From the 18 mutations in *walKR*, only one mutated position has been previously described (WalK-G223), which in previous studies resulted in an amino acid substitution at position 223 from glycine to aspartic acid (Howden et al., 2011; Hu et al., 2015; Vidaillac et al., 2013), and in our case to alanine. The alleles mutated strongly correlate with the ones found in clinical isolates, although most of the specific mutations are different, contributing to an expansion of the tolerance alleleome.

Vancomycin targets the cell wall, therefore, the finding that most of the key mutated genes under both TALE media conditions were related to cell wall biosynthesis was expected (i.e., *sgtB, prsA, walKRyycHI, vraSRT, pbp2, murJ_2, mprF*)(Hiramatsu, 2001; Jousselin et al., 2015; Kuroda et al., 2003; Łęski and Tomasz, 2005; Oku et al., 2004; Villanueva et al., 2018; Wang et al., 2001). For example, Pbp2 is the only bifunctional *S. aureus* penicillin-binding protein (transglycosylase and transpeptidase activities) (Goffin and Ghuysen, 1998; Murakami, 1994) and is involved in cell wall cross-linking. Pbp2 has also been associated with susceptibility to membrane and cell-wall targeting antibiotics (Łęski and Tomasz, 2005; Sieradzki and Tomasz, 1999). Besides the shared mutations, there were a number of media-specific mutations (Figure 3B). This environmental dependency was also evident from the expression of these key mutated genes (Figure 3C). For example, *vraG* and *mprF* genes that are mostly mutated in CA-MHB condition were highly expressed in vancomycin-tolerized strains in the same media, while there was no differential expression in RPMI+. On the other hand, some genes that seem to be mutated in a condition-specific manner presented a similar transcriptional profile in both (*e.g., sgtB* and *prsA*). Again, most of the key mutated genes were related to cell wall biosynthesis (e.g., *sgtB, prsA, walK, vraT, pbp2, mprF*). However, there were other key mutated genes associated with transcription (*rpoC*), transport (*pitA_1, vraG*), regulation (*perR, melR_2*), metabolism (*mqo2*), pathogenesis (*ebh_1*) and unknown function (*RS08780*).

Mutational analysis of TALE clones from CA-MHB revealed an instance of parallel evolution involving excision of the prophage ΦSa3 in three independent lineages. Large identical genomic deletions of 43,048 bp resulted from the excision of prophage ΦSa3, the most prevalent prophage family in *S. aureus (Xia and Wolz, 2014)*, which encodes for the immune evasion cluster (Goerke et al., 2009; Verkaik et al., 2011). This cluster harbors the immune modulators staphylokinase (Sak), staphylococcal complement inhibitor (SCIN), staphylococcal enterotoxin A (Sea), and chemotaxis inhibitory protein of *S. aureus* (CHIPS) (Read et al., 2018; van Wamel et al., 2006; Verkaik et al., 2011). Excision of the prophage results in the repair of the β-hemolysin gene (*hlb*) (Figure 4A) (Tran et al., 2019). Selective excision of prophage ΦSa3 has been reported, suggesting it acts as a molecular regulatory switch for β-hemolysin production (Tran et al., 2019). Testing of hemolytic activity of two of the three TALE strains and their starting strain counterparts confirmed one other characteristic phenotype observed in VISA strains, reduced hemolytic activity (Figure 4B). Since *hlb* encodes for a cold active hemolysin, after 24h incubation at 4 °C, it was possible to observe acquired hemolytic activity for the TALE strains which have excised the prophage ΦSa3 (Figure 4C), strains SVAM_A10 and SVAM_A12. In contrast, none of the RPMI+ media tolerized strains excised the prophage (Figure 3B). The expression of genes encoded in the prophage was analyzed in TALE strains derived from both media environments and it was observed that there was higher transcriptional activity of prophage genes in RPMI+ as compared to CA-MHB, for strains which retained the prophage genes (Figure 4D). This observation suggests an advantage in maintaining these prophage genes in RPMI+ upon vancomycin stress.

**Figure 4.**
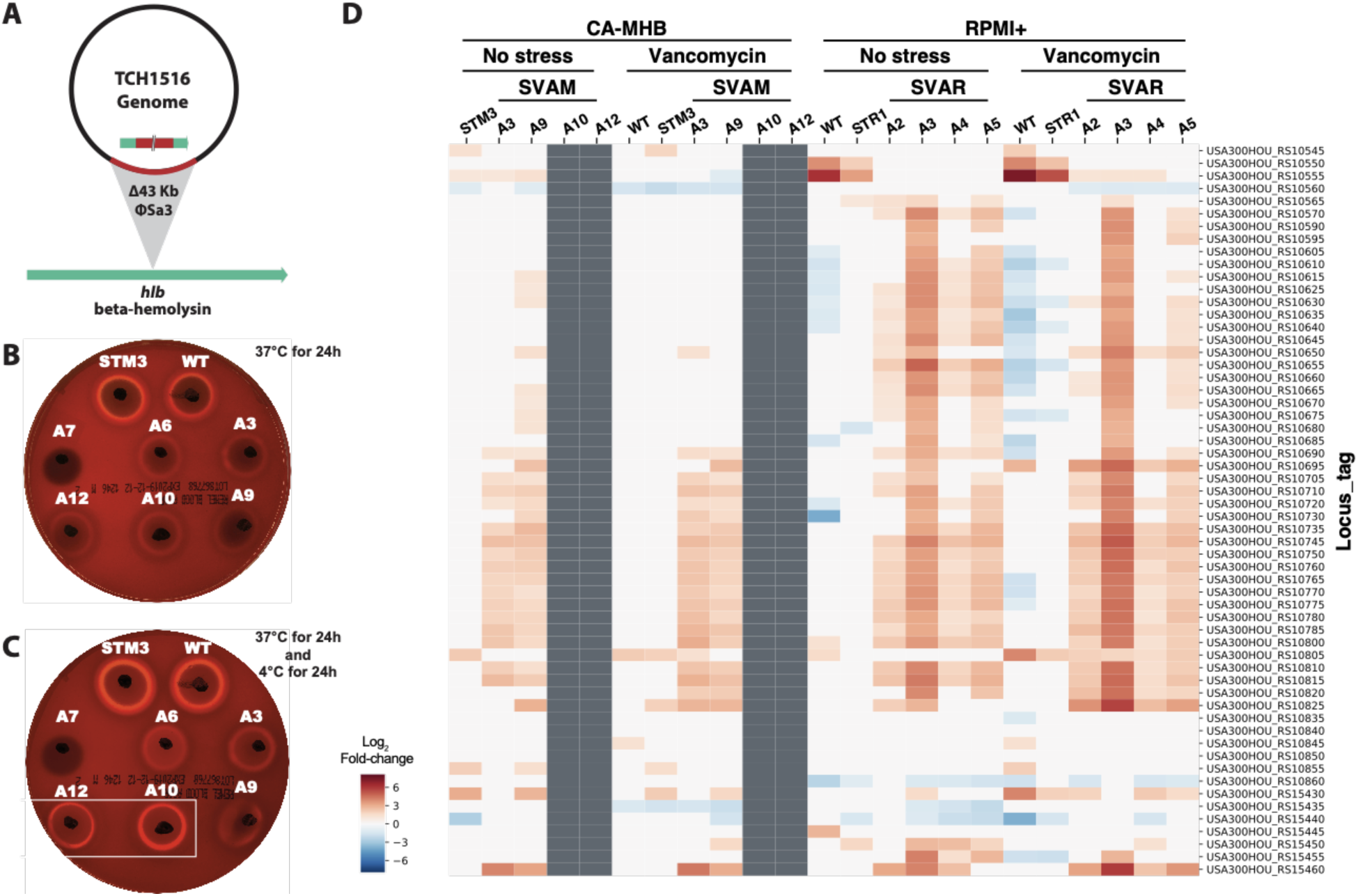
The excision of ΦSa3 prophage. (A) Schematic representation of the excision of the ΦSa3 prophage from the TCH1516 genome which leads to the repair of *hlb* gene, encoding for a β-hemolysin. (B) An image of a plate displaying hemolytic activity after 24 h incubation at 37 °C for vancomycin TALE strains in CA-MHB and their corresponding starting strains, wild-type (WT) TCH1516 and CA-MHB media-adapted STM3. (C) An image of the same plate in B displaying hemolytic activity following an additional 24 h incubation at 4 °C, to assess cold hemolytic activity of β-hemolysin. Increased hemolytic activity can be seen for strains SVAM_A10 and SVAM_A12 (boxed). (D) A heatmap of expression levels of genes encoded within the prophage ΦSa3. Grey indicates absence of the gene in a strain due to excision of the prophage. STM: *Staphylococcus aureus* adapted to CA-MHB. STR: *Staphylococcus aureus* adapted to RPMI+. SVAM: *Staphylococcus aureus* tolerized to vancomycin in CA-MHB. SVAR: *Staphylococcus aureus* tolerized to vancomycin in RPMI+.

### Broad scale impact of mutations in regulatory genes

The most commonly mutated genes in both media conditions during the TALE experiments were annotated with regulatory functions, with the *walRKyycHI* and *vraRST* operons being the most targeted (Figure 3B). Interestingly, the accessory genes (i.e., *yycH, yycI* and *vraT*) known to impact the activity of these regulators (Boyle-Vavra et al., 2013; Cameron et al., 2016), were some of the most often mutated. Mutations in regulatory genes tend to impact bacterial responses on a broad scale, which is difficult to assess solely from mutational data. Therefore, we performed RNAseq with and without vancomycin stress to understand how the mutations observed impacted the transcriptional profile of the evolved strains. Within the previously characterized WalR regulon (Delauné et al., 2012), there were several genes differentially regulated in the vancomycin TALE clones. In fact, both upregulation and downregulation was observed in several genes within this regulon (Figure 5A, and Supplementary Figure 3). Downregulation of the *spa* gene and lower hemolytic activity are a characteristic of VISA strains (Howden et al., 2010), which was also confirmed from the acquired transcriptional data (Figure 5A). The pyrimidine operon has been deemed important for growth in RPMI+ (Poudel et al., 2020) and was upregulated in strains evolved in this medium. The VraR regulon, responsible for the control of the cell wall stimulon, showed an overall upregulation in vancomycin-adapted strains (Figure 5B). The most upregulated genes, *vraX* and *cwrA*, are described to be part of both regulons (Figure 5), and have both been linked to cell wall stress response (Balibar et al., 2010; F. McAleese et al., 2006). Even though all the characterized vancomycin-tolerized strains presented a similar transcriptional rearrangement of these regulons, the analyzed strains carried distinct mutations in these regulatory genes (35 and 42 total different mutations in the *vra* and *wal* regulons, respectively; Supplementary Tables 3 and 4), suggesting that multiple mutational mechanisms can result in a similar transcriptional landscape. Overall, the TALE-derived mutations in these regulatory systems form a defined set of multiple unique mutations and resulted in significant rearrangements in expression levels of genes strongly associated with the observed tolerance phenotypes.

**Figure 5.**
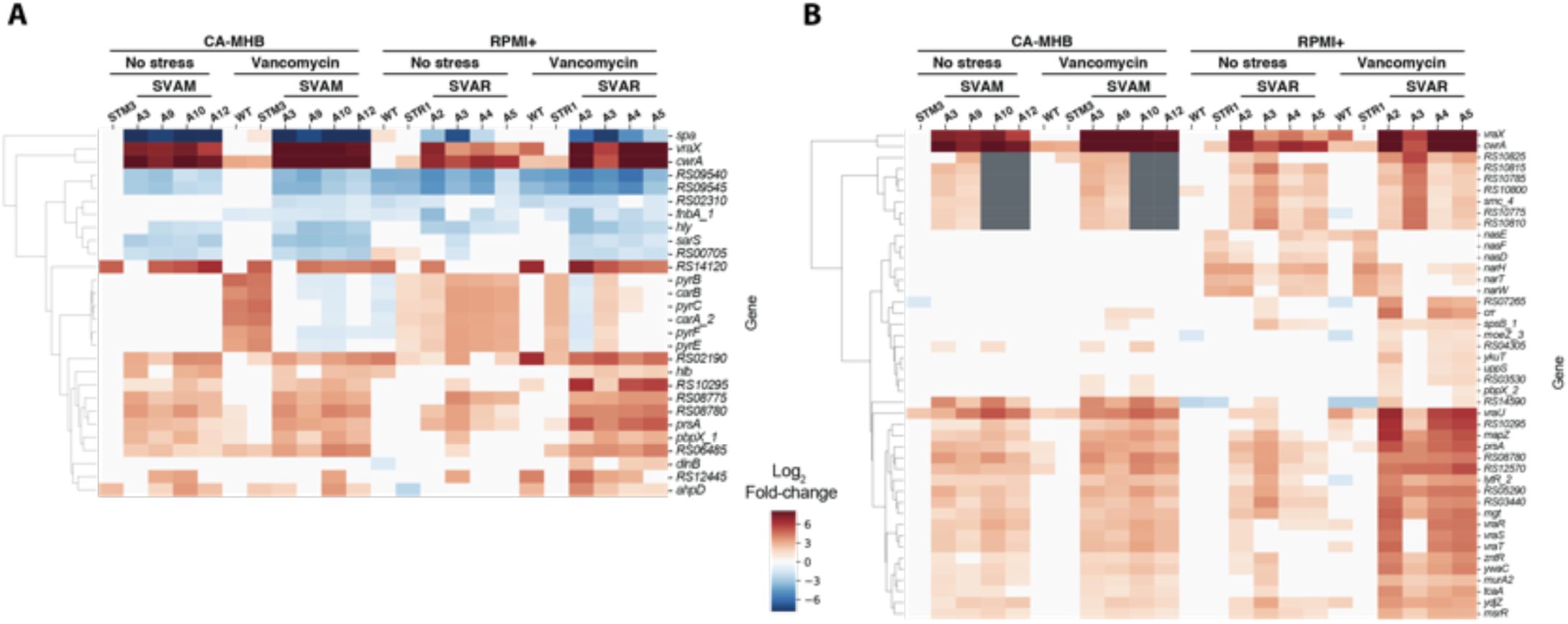
Rearranged transcriptional landscapes of operons associated with vancomycin tolerance in TALE strains. (A) A heatmap of expression levels of the genes in the WalR regulon (Delauné et al., 2012) displaying significant levels of differential expression, both up and down. (B) Expression of genes in the VraR cell wall stimulon (Boyle-Vavra et al., 2013). STM: *Staphylococcus aureus* adapted to CA-MHB. STR: *Staphylococcus aureus* adapted to RPMI+. SVAM: *Staphylococcus aureus* tolerized to vancomycin in CA-MHB. SVAR: *Staphylococcus aureus* tolerized to vancomycin in RPMI+.

### Elucidation of molecular mechanisms in vancomycin tolerance

Multiple mutations in *mprF* were observed in CA-MHB vancomycin-tolerized strains and were validated to display a significant decrease in negative cell surface charge. *mprF* encodes for a multi peptide resistance factor and has been associated with increased resistance to daptomycin and antimicrobial peptides (Ernst et al., 2018; Ernst and Peschel, 2019). Besides the previously described regulatory gene mutations, *mprF* was the most mutated gene in strains tolerized to vancomycin in CA-MHB. In the 12 CA-MHB-tolarized lineages, we identified seven previously reported mutations that lead to resistance through a MprF mechanism, and extended this knowledge base by identifying 10 new mutations (Figure6A). The MprF described resistance mechanism consists of decreasing the negative cell surface charge, and therefore repulsing cationic antimicrobial peptides and daptomycin (Ernst and Peschel, 2019). Since vancomycin is also a positively-charged peptide, one can speculate that this general mechanism would lead to increased vancomycin tolerance. In order to support this, we characterized the cell surface charge in tolerized mutants and its starting strain counterparts (Figure 6B). Tolerized mutants, with amino acid changes towards the C-terminal part of MprF, had a significant decrease in negative cell surface charge (Figure 6B and 6C). Strain SVAM_A3, with an amino acid change at position 50 (R50C), did not have a significant change in cell surface charge, which explains the observation of mutational hot-spots between positions 278 to 510 (Figure 6A). Thus, a similar mechanism of shifting the cell surface charge from more to less negatively-charged was validated for a subset of the *mprF* mutants uncovered from the TALE experimentation in this study.

**Figure 6.**
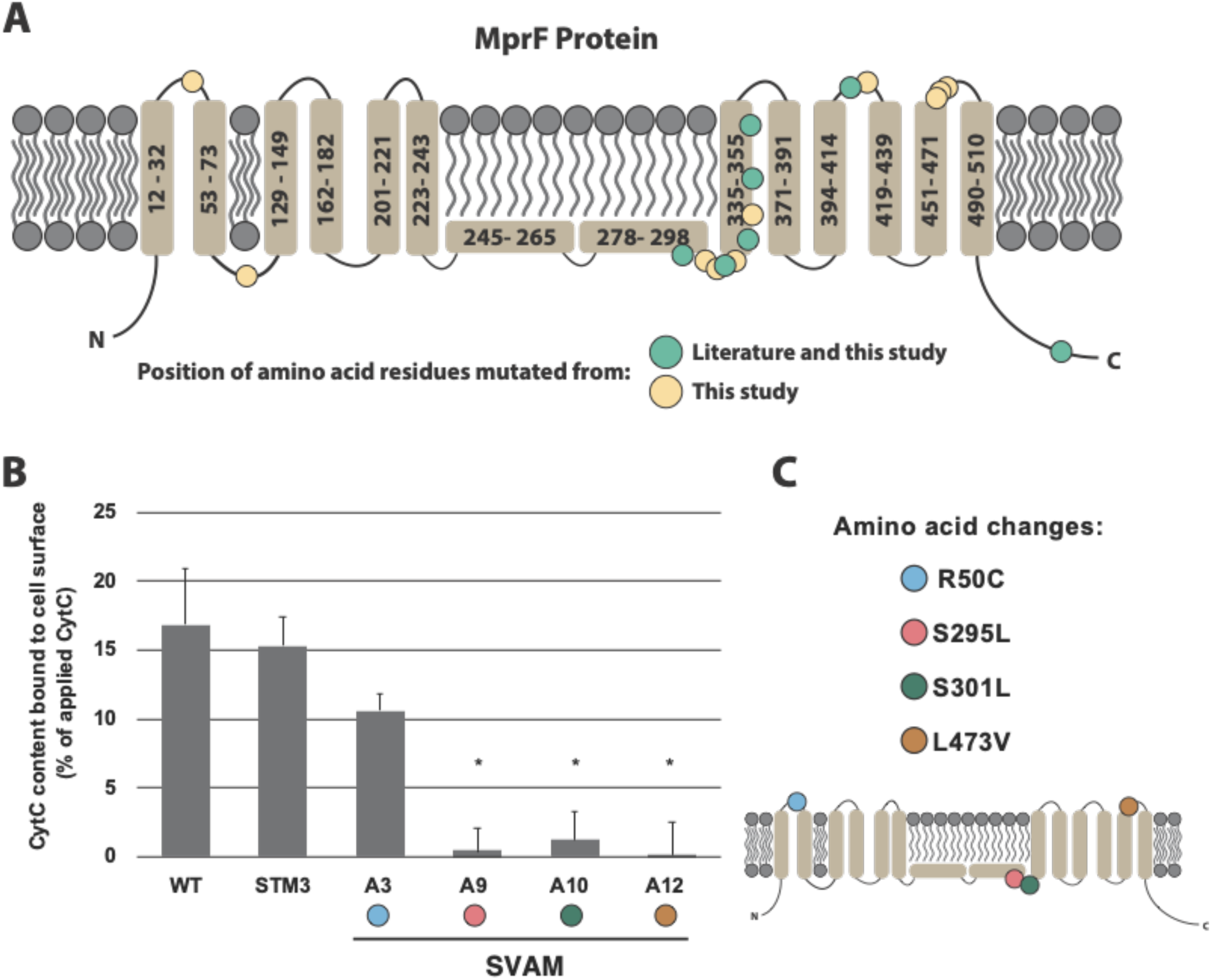
MprF mutations and their effect on cell surface charge. (A) Topology of the MprF membrane protein and mapping of mutations identified in this dataset (yellow and green circles), in comparison to previously reported ones (green circles) (Ernst and Peschel, 2019). (B) A graph quantifying the cell surface charge of strains tolerized to vancomycin, in comparison to their starting strain counterparts. Values that are significantly different (P ≤ 0.05) from the value for the respective starting strain (WT or STM3) by Student’s t-Test are indicated by an asterisk. (C) Identified locations of the specific amino acid substitutions observed in the tolerized strains and their position in the MprF structure. These mutations correspond to the strains tested in panel B. STM: *Staphylococcus aureus* adapted to CA-MHB. SVAM: *Staphylococcus aureus* tolerized to vancomycin in CA-MHB.

### Association of genetic targets with decreased susceptibility

Gene inactivation enables the assessment of a given gene’s role in tolerance. In order to understand the importance of the mutated and highly expressed genes in vancomycin tolerance, we relied on the available *S. aureus* Nebraska Transposon Mutant Library (Fey et al., 2013). We identified five genes required for tolerance and five genes impairing tolerance. A selection of mutants in key mutated genes or highly expressed genes was evaluated for their vancomycin susceptibility in both environmental conditions (Table 1). Interestingly, reduced vancomycin susceptibility was seen for all strains when testing was performed in RPMI+ compared to CA-MHB, in some instances with 2-fold or greater increases in vancomycin MIC in RPMI+. Media-specific susceptibility was observed for several mutants, similar to what was observed with the TALE-derived clones. Transposon inactivation of genes *yycI, sgtB, melR_2, stp*, and *lytM_2* resulted in a decreased susceptibility to vancomycin in RPMI+, but no difference in CA-MHB. On the other hand, *vraT, vraR, vraS, vraG* and *mprF* showed an increased susceptibility to vancomycin in CA-MHB, but no difference in RPMI+. The fact that gene inactivation can lead to decreased susceptibility is noteworthy as 57 (15.3%) and 76 (16.5%) of the total mutations identified lead to likely gene disruption in the RPMI+ and CA-MHB datasets, respectively. It is noteworthy that some of the gene disruptions identified in our datasets were in the same genes that showed here a higher MIC upon inactivation, specifically *yycI* and *stp* (in both media conditions), *melR_2* (only in RPMI+), and *lytM_2* (only in CA-MHB) (Supplementary Tables 3 and 4). These findings not only confirm the role of several genes (i.e. *yycI, sgtB, melR_2, stp, lytM_2, vraT, vraR, vraS, vraG* and *mprF*) in vancomycin tolerance, but also highlight the different roles these have in the development of resistance under varying environmental conditions.

**Table 1.** Minimum inhibitory concentration (MIC_90_) of *S. aureus* Nebraska Transposon Mutant Library (Fey et al., 2013) in both environmental conditions, CA-MHB and RPMI+. Strain ID refers to the NE identifier and the wild-type strain used in the generation of the transposon mutants (JE2). Gene indicates the gene that has been disrupted by the transposon.

| Strain ID | Gene | MIC<sub>90</sub> (µg/mL) |  |
| --- | --- | --- | --- |
|  |  | CA-MHB | RPMI+ |
| JE2 | - | 1 | 2 |
| NE1693 | *yycH* | 1 | 2 |
| NE1865 | *yycI* | 1 | 4 |
| NE596 | *sgtB* | 1 | 4 |
| NE1304 | *melR_2* | 1 | 4 |
| NE1919 | *stp* | 1 | 4 |
| NE641 | *lytM_2* | 1 | 4 |
| NE274 | *vraT* | 0.5 | 2 |
| NE554 | *vraR* | 0.5 | 1 - 2 |
| NE823 | *vraS* | 0.5 | 2 |
| NE70 | *vraG* | 0.25 - 0.5 | 2 |
| NE1360 | *mprF* | 0.25 - 0.5 | 2 |
| NE665 | *perR* | 0.5 - 1 | 2 |
| NE387 | *cwrA* | 1 | 2 |

## Discussion

The expanding problem of antibiotic resistant pathogens has been discussed for almost a century (Podolsky, 2018), but we remain in the infancy of understanding its complexity. In this study, we sought to understand changes in susceptibility, mutational mechanisms which enable such chances, and phenotypic responses of tolerant strains by analyzing growth screens, transcriptomics, and specific assays for evolved and mutated strains of MRSA under multiple media conditions. The main findings from this work are that (i) TALE successfully generated tolerant strains in both media types, although tolerance phenotypes translated across media types for CA-MHB TALE-derived strains, the inverse was not true for RPMI+; (ii) analysis of key mutational mechanisms revealed that numerous genetic allele variations can lead to similar transcriptional and phenotypic changes, especially in the CA-MHB condition; and (iii) TALE-derived strains shared similar properties to resistant clinical isolates in phenotype, mutation types, and gene expression.

There were a number of specific findings that provide context for the main conclusions of this study. These include, (i) the majority of TALE-derived strains tested maintained a tolerant phenotype after prolonged evolution under a no vancomycin stress condition; (ii) greater heterogeneity was found in the mutations observed in TALE-derived strains to become tolerant in CA-MHB than RPMI+, with a set of shared mutational mechanisms on the gene level from TALE-derived strains under both media conditions (enriched in cell wall related regulators), which resulted in major changes in expression for the operons they regulate even though the unique alleles differed; (iii) mutations in *mprF* are a key mechanism in CA-MHB, which decreased the overall negative cell surface charge and supposedly limited vancomycin access to the cell wall; (iv) similarly to clinically isolated strains, TALE-derived strains had lower hemolytic activity and reduced autolysis (Howden et al., 2010), mutations of the nature of pseudogenization (Mwangi et al., 2007), and similar transcriptional changes for virulence associated genes (e.g., *spa* (Howden et al., 2008; Fionnuala McAleese et al., 2006), *agr* (Mwangi et al., 2007; Sakoulas et al., 2002)); and lastly, (v) different genetic targets have enhancing or impairing roles in tolerance, depending on the environmental condition. Taken together, these findings provide specific information for a change in susceptibility for an important pathogen, an antibiotic commonly used to treat it, and media conditions relevant for human host infection and antimicrobial testing. Moreover, they provide context for the overall development of antibiotic resistance under multiple conditions and independent lineages that can be used to understand the issue as a whole.

The environmental conditions analyzed here were intended to represent both the *in vivo* environment simulating the condition of human infection, and the environment under which standard antimicrobial susceptibility is performed in the clinical laboratory. We showed that the evolutionary strategies adopted in each condition overlap on two major regulatory systems, VraR and WalR, which are responsible for the homeostasis of cell wall (Boyle-Vavra et al., 2013; Dubrac et al., 2007; McCallum et al., 2011; Villanueva et al., 2018) and had been previously linked to glycopeptide resistance (Howden et al., 2010; Hu et al., 2016; Kato et al., 2010). More interesting is the fact that besides such regulatory mutations, which seem to confer similar transcriptional rearrangements, other mutations were largely media-dependent, suggesting that the different media utilized restricted the evolutionary process differently. We have previously shown that growth of *S. aureus* in RPMI+ and CA-MHB lead to large transcriptional landscape changes (Poudel et al., 2020). In RPMI+, the buffering system used is bicarbonate, a ubiquitous buffer found in humans, which has been shown to potentiate the activity of several antibiotics by dissipating the proton motive force in bacteria (Ersoy et al., 2017; Farha et al., 2018; Kumaraswamy et al., 2016). This process might be one of the major reasons for the distinct evolutionary strategies observed under different media. For instance, this is likely the reason why we do not observe *mprF* mutations in the RPMI+ evolved strains, but we do in all the CA-MHB TALE lineages. MprF activity leads to the alteration of the cell surface charge (Ernst et al., 2018; Ernst and Peschel, 2019), but in the case of RPMI+ the membrane proton motive force is already compromised by the bicarbonate buffer system (Farha et al., 2018), making it an evolutionarily less viable solution towards vancomycin tolerance in this media. The tolerance mechanism through *mprF* mutation also explains the extended lag phase observed for all the strains tolerized in CA-MHB media, since an altered cell surface charge impacts bacterial cell division (Li et al., 2016; Strahl and Hamoen, 2010).

Even though both media used in vancomycin tolerizations resulted in highly tolerant vancomycin strains, in the case of strains evolved in RPMI+, this phenotype did not translate to CA-MHB media. Media dependent susceptibilities have been previously reported (Ersoy et al., 2017; Farha et al., 2018; Kumaraswamy et al., 2016; Lin et al., 2015). Here we showed that the evolution of resistance in physiological conditions is not phenotypically revealed in clinical susceptibility testing (Figure 2B). The fact that decreased vancomycin susceptibility acquired in RPMI+ simulates the selective conditions under which it evolves in patients receiving vancomycin (McKee and Komarova, 2017), and that these changes are not detected in the CA-MHB utilized laboratory testing essentially shows that the clinical laboratory is blind to the clinically relevant reduction in vancomycin susceptibility that evolves in *S. aureus*. This may explain the poor clinical efficacy of vancomycin even against *S. aureus* isolates that the laboratory designates as susceptible. Clinical experience is abundant with patients with MRSA bacteremia by organisms fully susceptible to vancomycin, yet fail to clear their infection despite adequate dosing and the lack of surgical solutions. The clinical laboratory shortcomings in detecting vancomycin resistance in *S. aureus* may be one explanation for the fact that vancomycin is unique among anti-staphylococcal antibiotics where resistance took decades to emerge (at least according to the laboratory). Resistance to every anti-staphylococcal antibiotic has emerged just a few years after the clinical introduction of that antibiotic, yet for vancomycin, which was introduced into clinical practice in 1958, VISA was not described until 1997 (Levine, 2006). Examining these data at an even higher level shows that every mutant from the Nebraska library demonstrated a higher vancomycin MIC in RPMI+ compared to CA-MHB. Given that the area under the curve (AUC)/MIC ratio is the pharmacokinetic target reflective of vancomycin activity suggests considerably weaker activity of vancomycin *in vivo* than in clinical laboratory conditions would indicate (Giuliano et al., 2010). Indeed, the well-described clinical-microbiological discordance of vancomycin with regards to *S. aureus* explained by our findings supports serious re-examination of how antimicrobial susceptibility paradigms can be made more clinically relevant (Ersoy et al., 2017).

During the many years that antibiotic resistance has been studied, associations have been made between specific alleles and decreased antibiotic susceptibility (Cameron et al., 2012; Howden et al., 2011, 2010; Hu et al., 2015, 2016; Ishii et al., 2015; Mwangi et al., 2007; Vidaillac et al., 2013). Numerous surveys have been conducted using PCR and sequencing to determine the likelihood of a given strain to be less susceptible to a given antibiotic (Costa et al., 2018; Kato et al., 2010; Sabat et al., 2018; Shore et al., 2010). Here we showed that limiting the analysis to a handful of genes can be misleading and that many allele variants can result in the same outcome. We identified several mutations in alleles previously associated with decreased susceptibility, with most of the mutations being new variants. A high-throughput approach utilizing ALE was an efficient way to sample the evolutionary pathways available for the development of antibiotic resistance, while expanding on the knowledge of allelic variation responsible for such phenotypes. Examples provided here are the mutations in the regulatory systems (i.e., *vraSRT* and *walKRyycHI*) and in *mprF*. We showed that different allele variants in regulatory genes can impact the transcriptional landscape similarly. We have largely expanded the knowledge on *mprF* allele variants that result in altered cell surface charge that might lead to decreased vancomycin susceptibility and bridge the knowledge gap between vancomycin and peptide antibiotic cross-resistance.

Pseudogenization is another type of genetic variance we believe merits attention. We previously showed that *S. aureus* restored pseudogenes in order to overcome metabolic limitations (Machado et al., 2019). In this dataset, approximately 15% of all the mutations led to pseudogenization, which can translate into decreased susceptibility, as demonstrated with the transposon mutants (Table 1). This strengthens the hypothesis that *S. aureus* can use this pseudogenization mechanism to adapt to distinct environments, including the development of antibiotic resistance. A study looking at the adaptive evolution of *S. aureus* during chronic endobronchial infection of a cystic fibrosis patient during 26 months identified 391 mutations (comparable to our datasets) with none of the mutations predicted to result in pseudogene formation (McAdam et al., 2011). The rates at which pseudogenization occurs and reverts, requires further experimental evidence in order to support this method as a common evolutionary strategy in *S. aureus*.

In conclusion, the application of ALE to develop *S. aureus* strains tolerized to vancomycin was successful and links can be drawn between TALE-derived strains and clinical isolates. Susceptibility was not only decreased for the targeted antibiotic, but also reproduced phenotypes similar to the ones previously reported for clinical strains with decreased vancomycin susceptibility (Howden et al., 2010). Furthermore, it allowed us to understand evolutionary strategies and constraints in two clinically relevant media environments while expanding our knowledge on the diversity of alleles contributing to the vancomycin tolerant phenotype. These findings allow a better understanding of the evolution of antibiotic resistance and provide new information valuable for the epidemiological surveillance and control of *S. aureus* resistance in clinical environments. Most importantly, these data call into question the clinical reliability of *S. aureus* vancomycin susceptibility testing as it is currently performed in the clinical laboratory by providing a deeper understanding of why *S. aureus* resistance to vancomycin is rare in the laboratory yet vancomycin treatment failure is common in clinical practice.

## Material and Methods

### Tolerization Adaptive Laboratory Evolution (TALE)

TALE was performed as previously described (Mohamed et al., 2017), with variations as noted. Four replicates of each starting strain (wild type and two media-adapted strains per media type) were inoculated from independent colonies on LB-agar plates. Cultures were grown in 15 ml working volume tubes which were heated to 37°C and were aerobically stirred at 1100 rpm. Periodically, optical density readings at a 600 nanometer wavelength (OD_600_) were taken for each culture with a Tecan Sunrise reader plate, until the OD_600_ reached approximately 0.6 (approximately equivalent to an OD_600_ of 1 on a cm path length reader). At that time, 150 µl of the culture was passed to a fresh tube, to prevent the cells from reaching stationary phase. The passage volume was adjusted dynamically based on the actual OD_600_ at the time of passage, to keep the number of cells passed consistent. Additionally, if the culture had grown for several consecutive flasks (∼3 flasks), the vancomycin concentration in the next tube was increased. This stepwise increase began at 20% of the starting concentration but augmented over the course of the experiment. Growth rates were estimated for each tube by linear regression of the natural log of the optical density vs. time. Periodically throughout the experiment, culture aliquots were taken for long term storage at −80°C by mixing 800 µL of 50% glycerol with 800 µL of culture.

### Whole genome re-sequencing

DNA sequencing was performed on clones and populations throughout the evolution, covering two or three timepoints of the evolution. Total genomic DNA was sampled from an overnight culture and extracted using a KingFisher Flex Purification system previously validated for the high throughput platform mentioned below (Marotz et al., 2017). Sequencing libraries were prepared using a miniaturized version of the Kapa HyperPlus Illumina-compatible library prep kit (Kapa Biosystems). DNA extracts were normalized to 5 ng total input per sample using an Echo 550 acoustic liquid handling robot (Labcyte Inc), and 1/10 scale enzymatic fragmentation, end-repair, and adapter-ligation reactions carried out using a Mosquito HTS liquid-handling robot (TTP Labtech Inc). Sequencing adapters were based on the iTru protocol (Glenn et al., 2019), in which short universal adapter stubs are ligated first and then sample-specific barcoded sequences added in a subsequent PCR step. Amplified and barcoded libraries were then quantified using a PicoGreen assay and pooled in approximately equimolar ratios before being sequenced on an Illumina HiSeq 4000 instrument.

The obtained sequencing reads were trimmed and filtered using AfterQC software, version 0.9.6 (Chen et al., 2017). Re-sequencing analysis for mutation identification was performed using the breseq bioinformatics pipeline (Deatherage and Barrick, 2014), version 0.31.1 and the *S. aureus* TCH1516 reference genome (GCA_000017085.1), reannotated using PATRIC (Brettin et al., 2015). ALEdb was used for mutation analysis (Phaneuf et al., 2018).

### Minimum inhibitory concentration

Strains were pre-cultured in the corresponding media (CA-MHB or RPMI+10%LB) for ∼5h, and then inoculated to a final OD of 0.002 in media with or without vancomycin (Sigma). Growth was measured by following OD_600_ in a Bioscreen C Reader system with 150 μL per well. MIC_90_ was determined at 17 h post incubation. The experiments were done in biological triplicates.

### Estimation of growth parameters

Growth parameters were estimated as previously described (Anand et al., 2020). Briefly, lag phase was estimated by fitting the Baranyi growth model (Baranyi and Roberts, 1994) using nonlinear regression in R. A sensitivity analysis was run to exclude data points beyond a specific time threshold T, to avoid skewing the estimated parameters as a result of a possible cell death phase, secondary growth phase or noise. The sensitivity analysis ensured that the lag phase, exponential phase and stationary phase only are taken into account in the fitting process, because all other growth/death phases are not explicitly modeled in Baranyi’s equation. Anova was run in R using aov() to test the null hypothesis that there is no difference in lag phase duration between pre-evolved strains (WT, STM2, STM3, STR1 and STR4) and vancomycin adapted strains.

### Resistance phenotype stability

Following TALE adaptation to vancomycin, a subset of the TALE strains (6 from each media) were further evolved in duplicate in the respective media for 21.79 ± 2.08 passages, 9.41×10^11^ ± 9.84×10^10^ CCDs. The final populations were then evaluated for their vancomycin susceptibility as described above.

### Transcriptomics

Eight TALE strains were selected for transcriptional analysis, along with their pre-evolved counterparts (wild type and one media-adapted strain per medium). Total RNA was sampled from biological duplicate cultures. The strains were grown in each respective media, with and without vancomycin (at 1/2 MIC). At OD 0.2, cultures without vancomycin were harvested. At the same OD, cultures were treated with 0.5x MIC of vancomycin, and harvested for RNA 30min after. Harvesting of the cells consisted in mixing of 3 mL of culture with two volumes of Qiagen RNA-protect Bacteria Reagent (6 ml), vortexed for 5 s, incubated at room temperature for 5 min and immediately centrifuged for 10 min at 17,500 r.p.m.. The supernatant was decanted, and the cell pellet was stored at −80 °C. Total RNA was isolated using the Quick RNA Fungal/Bacterial Microprep (Zymo Research), following vendor procedures, including an on-column DNase treatment. RNA quality and purity was assessed using Nanodrop and Bioanalyzer RNA nano chip. Ribosomal RNA was removed from total RNA preparations using RNaseH. Then secondary structures in the ribosomal RNA were removed by heating to 90 degrees for 1 second, a set of 32-mer DNA oligo probes complementary to the 5S, 16S, and 23S subunits and spaced 9 bases apart were then annealed at 65 degrees followed by digestion with Hybridase (Lucigen), a thermostable RNAseH. The enzyme was added at 65 °C, the reaction incubated for 20 minutes at that temperature, then heated again to 90 °C for 1 second to remove remaining secondary structures, and returned to 65 °C for 10 minutes. The reaction was quickly quenched by the addition of guanidine thiocyanate while still at 65 °C before purifying the mRNA with a Zymo Research RNA Clean and Concentrator kit using their 200 nt cutoff protocol. Carryover oligos were removed with a DNAse I digestion which started at room temperature and gradually increased to 42 °C over a half hour. This was followed up with another column purification as stated above. The remaining RNA was used to build a cDNA library for sequencing using a KAPA Stranded RNA-seq Library Preparation Kit. The generated cDNA libraries were sent for Illumina sequencing on a HiSeq 4000.

The phred quality scores for the Illumina sequencing reads were generated using FastQC package (Andrews and Others, 2010). Bowtie2 was used to align the raw reads to TCH1516 genome (GCA_000017085.1) and to calculate alignment percentage (Langmead and Salzberg, 2012). The aligned reads were then normalized to transcripts per million (TPM) with DESeq2 (Love et al., 2014). The final expression values were log-transformed log_2_[TPM + 1] for visualization and analysis.

### Hemolysin production

Analysis of hemolysin production was performed by spotting 10 μL of a OD_600_ culture of 1 grown in CA-MHB onto 5% sheep blood agar plates. Plates were incubated at 37 °C for 24 h, followed by a 4 °C incubation for another 24 h. Hemolysin production was monitored after both incubations by observing the appearance of a clear halo.

### Surface charge

Quantification of the relative cell surface charge was performed using a cytochrome C (Sigma) binding assay, as previously described (Peschel et al., 1999). Briefly, cells were grown in CA-MHB until an OD_600_ of ∼2, washed twice with MOPS buffer (20 mM, pH 7) and finally resuspended to an OD_600_ of 5. These were incubated for 10 min with 0.5 mg/ml (cytochrome c), which was subsequently removed by centrifugation. The amount of cytochrome C was spectrophotometrically quantified at 530nm. The amount of cytochrome C bound to the cells can be used as a proxy for the cell surface charge. The experiments were done in biological triplicates.

### Autolysis assay

Evaluation of autolysis was performed using the Triton X-100-induced autolysis assay. Cells were grown to an OD_600_ of 1, washed twice with PBS buffer and resuspended in PBS buffer containing 0.05 % Triton X-100. Cell suspensions were incubated at 37 °C and autolytic activity was measured by monitoring the OD_600_ every hour using Tecan Infinite 200 Pro microplate reader. The experiments were done in biological triplicates.

### Accession number(s)

Newly determined DNA sequence data were deposited in the NCBI database under BioProject PRJNA521551, accession numbers SRR8552163 to SRR8552250. All RNA-seq data have been deposited to the GEO database (record GSE149213) and Short Read Archive (SRA), RNA-seq data accession numbers SRX8164260 to SRX8164307.

## Acknowledgements

This research was supported by NIH NIAID grant (U01-AI124316).

## Supplemental material

**Table S1 -** Minimum inhibitory concentration (MIC_90_) of *S. aureus* starting strains and TALE-derived strains in the same environmental conditions used for tolerance evolution, CA-MHB or RPMI+. Post-TALE refers to the MIC_90_of the population after 21.79 ± 2.08 passages (9.41×10^11^ ± 9.84×10^10^ CCDs) in the media used for evolution, without vancomycin. The post-TALE evolution was performed in duplicate for each of the end-point clones.

**Figure S1 –** Characteristics of vancomycin TALE strains. (A) An image of a plate displaying hemolytic activity after 24 h incubation at 37 °C for starting strains and vancomycin TALE strains (B) Autolysis of whole cells of *S. aureus*. Mid-exponential-phase cultures were resuspended in 0.05 M Tris-HCl (pH 7.2) containing 0.05% Triton X-100 and were incubated at 30°C. Absorbance was measured every hour and percentage to initial absorbance calculated.

**Table S2 –** ANOVA statistical analysis of the lag-phase duration of starting strains and evolved strains in the same media as that used for tolerization.

**Figure S2 –** Venn diagram of the key mutated genes (≥ 2 instances) in the two utilized media conditions (i.e., CA-MHB and RPMI+).

**Table S3 -** Mutations identified for TALE-derived strains after tolerization to vancomycin in CA-MHB. Nomenclature example A1 F29 I1 R1 = ALE 1 Flask 29 Isolate (I1=clone, I0=population) Replicate 1.

**Table S4 -** Mutations identified for TALE-derived strains after tolerization to vancomycin in RPMI+. Nomenclature example A1 F29 I1 R1 = ALE 1 Flask 29 Isolate (I1=clone, I0=population) Replicate 1.

---

**Table S1.** Minimum inhibitory concentration (MIC_90_) of *S. aureus* starting strains and TALE-derived strains in the same environmental conditions used for tolerance evolution, CA-MHB or RPMI+. Post-TALE refers to the MIC_90_of the population after 21.79 ± 2.08 passages (9.41×10^11^ ± 9.84×10^10^ CCDs) in the media used for evolution, without vancomycin. The post-TALE evolution was performed in duplicate for each of the end-point clones.

| Strain | MIC<sub>90</sub> (µg/mL) in CA-MHB |  | Strain | MIC<sub>90</sub> (µg/mL) in RPMI+ |  |
| --- | --- | --- | --- | --- | --- |
|  | Starting strain / TALE | post-TALE |  | Starting strain / TALE | post-TALE |
| WT | 1 | - | WT | 2 | - |
| SVAM_A2 | 4 | ND | SVAR_A1 | 4 | ND |
| SVAM_A3 | 8 | 4 / 4 | SVAR_A2 | 8-16 | 8 / 8 |
| STM2 | 1 | - | SVAR_A3 | 16 | 16 / 16 |
| SVAM_A6 | 4-8 | 2-4 / 4 | SVAR_A10 | 8 | ND |
| SVAM_A7 | 8 | 4 / 4 | STR1 | 2 | - |
| SVAM_A8 | 4 | ND | SVAR_A4 | 8-16 | 8 / 8 |
| STM3 | 1 | - | SVAR_A5 | 8-16 | 8-16 / 8 |
| SVAM_A9 | 8 | 4 / 4 | SVAR_A11 | 8 | ND |
| SVAM_A10 | 8 | 2 / 2 | STR4 | 4 | - |
| SVAM_A11 | 4 | ND | SVAR_A7 | 8-16 | 8 -16 / 16 |
| SVAM_A12 | 8 | 4 / 4 | SVAR_A8 | 8 | ND |
|  |  |  | SVAR_A12 | 8 | 8 / 8 |

**Figure S1.**
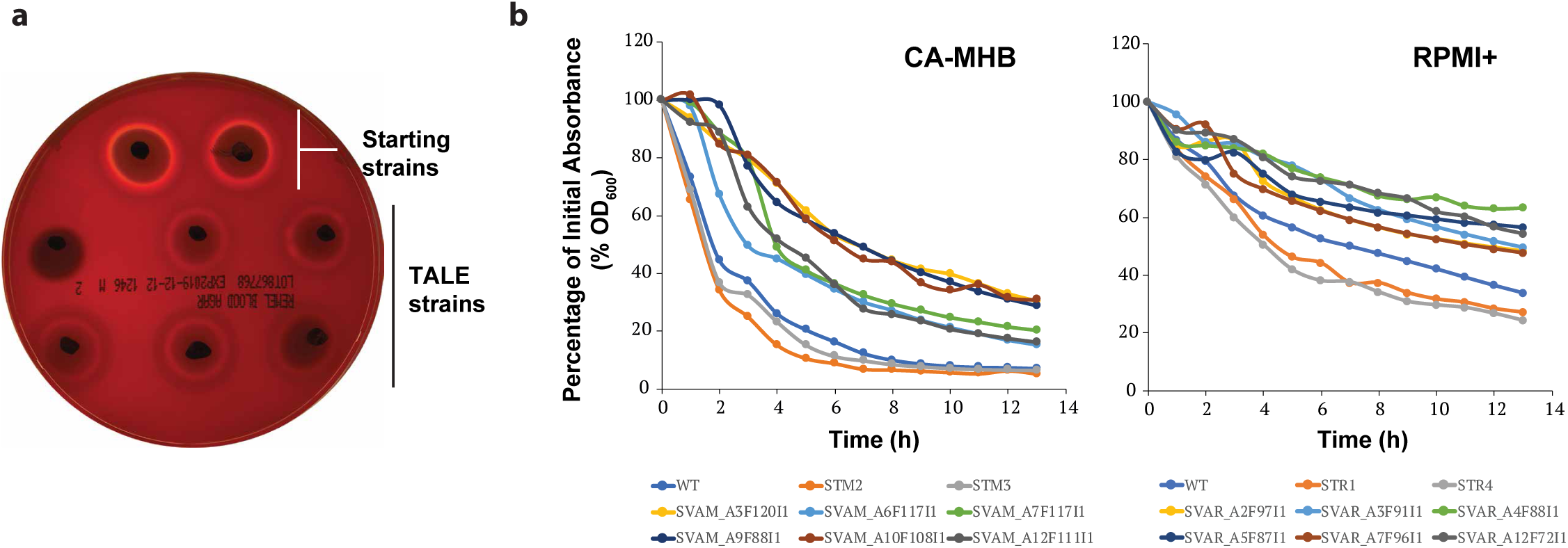
Characteristics of vancomycin TALE strains. (A) An image of a plate displaying hemolytic activity after 24 h incubation at 37 °C for starting strains and vancomycin TALE strains (B) Autolysis of whole cells of *S. aureus*. Mid-exponential-phase cultures were resuspended in 0.05 M Tris-HCl (pH 7.2) containing 0.05% Triton X-100 and were incubated at 30°C. Absorbance was measured every hour and percentage to initial absorbance calculated.

**Table S2.** ANOVA statistical analysis of the lag-phase duration of starting strains and evolved strains in the same media as that used for tolerization.

| vancomycin.ug.ml | Starting_strain | F.value | Pr..F. |
| --- | --- | --- | --- |
| 0 | STM2 | 4.37324747923248 | 0.062998644983095 |
| 0 | STM3 | 8.21404843584888 | 0.0132439301947123 |
| 0 | STR1 | 55.9776079042055 | 2.10636199647478E-05 |
| 0 | STR4 | 18.2400821654054 | 0.00163482854999148 |
| 0.5 | STM2 | 180.845045483513 | 9.94033084285709E-08 |
| 0.5 | STM3 | 163.726248981813 | 9.6409038322023E-09 |
| 0.5 | STR1 | 355.069596153815 | 3.83918455769995E-09 |
| 0.5 | STR4 | 198.53651002671 | 6.36874197639287E-08 |
| 0 | WT(MHB) | 533.791403846887 | 7.21720540302604E-08 |
| 0 | WT(RPMI) | 11.7094080873006 | 0.00652657253351374 |
| 0.5 | WT(MHB) | 499.257469737133 | 9.09544723581264E-08 |
| 0.5 | WT(RPMI) | 5.55414886661091 | 0.0401771670140192 |

**Figure S2.**
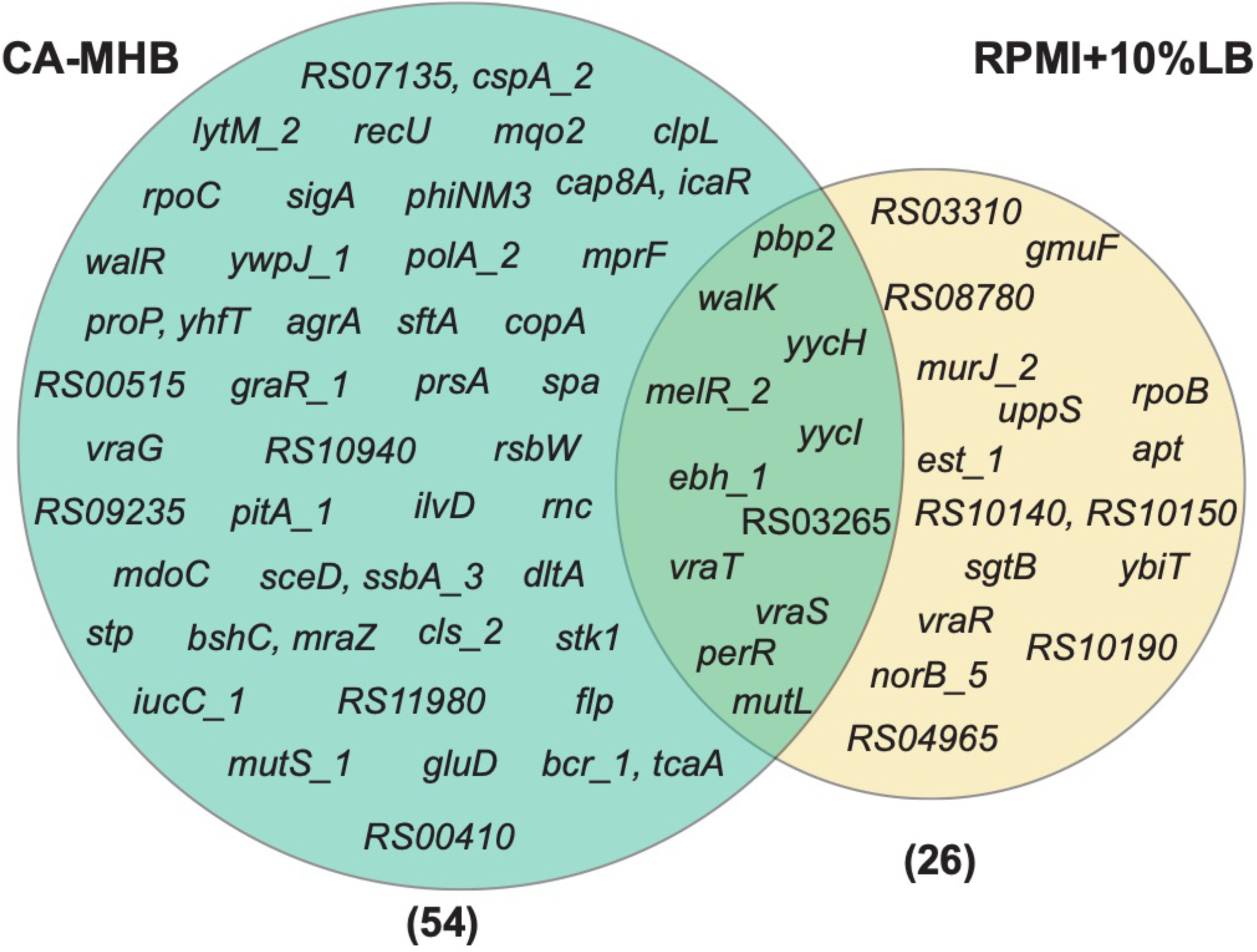
Venn diagram of the key mutated genes (≥ 2 instances) in the two utilized media conditions (i.e., CA-MHB and RPMI+).

